# Cell type-specific assessment of cholesterol distribution in models of neurodevelopmental disorders

**DOI:** 10.1101/2022.11.16.516849

**Authors:** Charlotte Czernecki, Shirley Dixit, Isabelle Riezman, Sabrina Innocenti, Caroline Bornmann, Frank W. Pfrieger, Howard Riezman, Peter Scheiffele

## Abstract

Most nervous system disorders manifest through alterations in neuronal signaling based on abnormalities in neuronal excitability, synaptic transmission, and cell survival. However, such neuronal phenotypes are frequently accompanied – or even caused – by metabolic dysfunctions in neuronal or non-neuronal cells. The tight packing and highly heterogenous properties of neural, glial and vascular cell types pose significant challenges to dissecting metabolic aspects of brain disorders. Perturbed cholesterol homeostasis has recently emerged as key parameter associated with sub-sets of neurodevelopmental disorders. However, approaches for tracking and visualizing endogenous cholesterol distribution in the brain have limited capability of resolving cell type-specific differences. We here develop tools for genetically-encoded sensors that report on cholesterol distribution in the mouse brain with cellular resolution. We apply these probes to examine sub-cellular cholesterol accumulation in two genetic mouse models of neurodevelopmental disorders, *Npc1* and *Ptchd1* knock-out mice. While both genes encode proteins with sterol-sensing domains that have been implicated in cholesterol transport, we uncover highly selective and cell type-specific phenotypes in cholesterol homeostasis. The tools established in this work should facilitate probing sub-cellular cholesterol distribution in complex tissues like the mammalian brain and enable capturing cell type-specific alterations in cholesterol flow between cells in models of brain disorders.

## Introduction

Neurodevelopmental conditions, including autism, are highly heterogeneous in their severity, phenotypic characteristics, and underlying mechanisms. This heterogeneity arises from diverse genetic, metabolic, and environmental factors that contribute to the etiology of the conditions. Consequently, therapies will need to be tailored specifically to the afflicted individuals, highlighting the importance of gaining insights into the origin and pathophysiology of these complex disorders ^1-3^. Previous work identified alterations in specific synaptic neurotransmitter receptor systems, neuromodulators, intracellular signaling pathways, and transcriptional regulators that arise from autism-associated mutations and that resemble candidate targets for interventions ^1, 4-9^. Lipid homeostasis is a comparably less deeply explored process that – when perturbed – increases the likelihood of neurodevelopmental conditions, including autism. There is a high incidence of autistic spectrum features in individuals with Smith-Lemli-Opitz syndrome (SLOS) with more than half of individuals meeting diagnostic criteria for autism ^10, 11^. SLOS arises from loss-of-function mutations in the 7-dehydrocholesterol reductase (DHCR7), an enzyme that catalyzes the conversion of 7-dehydrocholesterol to cholesterol. Metabolically, SLOS is characterized by cholesterol insufficiency and accumulation of its dehydrocholesterol precursors. Notably, alterations in cholesterol homeostasis are more widely associated with neurodevelopmental disorders (NDDs). Population-level differences in blood lipid profiles (LDL, total cholesterol, and triglycerides) were reported between individuals with ASD and matched controls ^12, 13^. Moreover, cholesterol alterations have been implicated in Fragile X Syndrome ^14^ and Rett’s Syndrome ^15^, two monogenic syndromes where va substantial fraction of individuals meets diagnostic criteria for autism. Finally, Niemann-Pick type C disease is a lysosomal storage disorder characterized by intracellular accumulation of unesterified cholesterol and other lipids and progressive neurodegeneration. Neurodevelopmental assessment of children with NPC1 reported a high incidence of developmental delay ^16^ that can appear before the onset of neurologic symptoms ^17^. Overall, defects in cholesterol homeostasis were hypothesized to play an important role in a sub-group of neurodevelopmental conditions and to contribute to the etiology of autism ^18^.

Blood lipid profiles are routinely analyzed in clinical practice, whereas detecting altered cholesterol homeostasis in the brain is complicated by the localized synthesis and transport of cholesterol across sub-cellular compartments and cell types. Cholesterol in the brain derives from local *de novo* synthesis and subsequent intercellular distribution ^19^. Neuronal cells exhibit very low level of cholesterol synthesis. Instead, cholesterol synthesized in astrocytes is released in form of lipoprotein particles which are internalized by neurons through receptor-mediated endocytosis ^20^. Considering that lipoproteins are only poorly transported across the blood-brain barrier, measurement of peripheral cholesterol provides little information about brain cholesterol levels. Moreover, measurements in brain extracts cannot inform about cholesterol distribution in specific brain cells such as neuronal, astrocytic, microglial cells, and oligodendrocyte-derived myelin membranes. Thus, new approaches are needed to probe cholesterol homeostasis in the brain.

The development of genetically-encoded cholesterol sensors provided a major advance in dissecting cellular distribution and dynamics of this essential membrane component *in vitro*. These probes are based on domain 4 (D4) of the bacterial toxin perfringolysin O which binds to accessible pools of cholesterol in cellular membranes ^21-23^. D4H derivates, such as the YDA variant, exhibit a range of minimum cholesterol concentrations required for binding. Thus, D4H and YDA are recruited to cellular membranes containing 20%mol and 2%mol cholesterol, respectively ^24^. These probes provided important insights into cholesterol dynamics associated with synaptic plasticity ^25, 26^. However, their use has been largely limited to cell culture systems as the probes exhibit fluorescent signals regardless of cholesterol binding which complicates distinguishing differential probe expression levels from differential cholesterol levels across cell types. Here, we further developed these probes to assess cholesterol distribution in the mouse brain. We generated a set of ratiometric cholesterol sensors to probe cholesterol distribution in a cell type-specific manner. Finally, we demonstrate the utility of these probes for the quantitative assessment of cell type-specific alterations of cholesterol in mouse models of NDDs.

## Materials and Methods

### Mice and animal experimentation

All animal experiments were approved by the Cantonal Veterinary Office of Basel-Stadt (Switzerland), and performed in accordance with Swiss laws. The following transgenic mouse lines were used in this study: *Npc1*^*KO*^ (BALB/cNctr-Npc1<m1N>/J, The Jackson Laboratory, strain #003092), *Ptchd1*^*KO* 27^ (B6.Cg-Ptchd1tm1d(KOMP)IcsOrl/Mmucd, MMRRC, stock n° 043797-UCD), PV-cre (B6;129P2-Pvalbtm1(cre)Arbr/J, The Jackson Laboratory, strain #008069). For all experiments, mice were strain-matched, age-matched, and wild-type littermates were used as controls. For the *Ptchd1*^*KO*^ line, only males were used in experiments, because the mouse and human *Ptchd1* genes are X-chromosomal.

### Viral and plasmid vectors

cDNAs for genetically-encoded mCherry-D4H/YDA cholesterol probes were kindly provided by Dr. Sophie Martin (UNIL, Lausanne, Switzerland) (pSM2056_Pact_mCherry_D4H-Tdh2). The mCherry-D4H/YDA coding sequence was fused to a eGFP and T2A sequence and inserted in a adenoviral backbone under control of the human Synapsin promoter (hSyn). As cholesterol-independent membrane probe, a myristoylated/palmitoylated-mCherry was used. For astrocyte-specific expression, the hSyn promoter was replaced by a minimal GFAP-promoter (GfaABC1D) ^28^. Viruses were generated in HEK293T cells using standard protocols with AAV9 capsid. Viral preparations were concentrated in 100K Millipore Amicon columns at 4°C. Samples were then suspended in PBS, aliquoted and stored at −80°C. Viral titers were determined by qPCR and were >10^13^ particles/mL.

### GeneTrek analysis

A curated list of sterol-sensing domain (SSD) proteins and sterol metabolism genes was created using KEGG pathways and HGNC nomenclature (Supplementary table 1), and entered in the GeneTrek search tool. GeneTrek is used to explore association of human genes with several neurodevelopmental disorders (NDD) including autism-spectrum disorders (ASD), intellectual disability (ID) and others ^29^.

### Cell culture experiments

HEK293T cells were grown on 10 mm glass coverslips in 24-well plates in DMEM medium (Thermo Fisher, cat n°10566016) supplemented with fetal bovine serum (10%) and penicillin/streptomycin (1%). 24h after plating, cells were transfected with 800 ng of plasmid vectors (50 ng / 750 ng or 200 ng / 600 ng of pAAV-Syn-eGFP-2A-mCherry_D4H/YDA or pAAV-Syn-Myr_mCherry-2A-eGFP and pAAV-iCre control respectively) using the FuGene6 transfection protocol (Roche, cat n°11 844 443 001) at a 3:1 DNA:FuGene ratio. At 24h post-transfection, cells were fixed with PFA 4% in PBS 1X (Electron Microscopy Sciences, cat n°15700) for 10 min, washed twice with PBS 1X, stained with DAPI 1/10000 and finally washed with PBS. Coverslips were subsequently mounted on glass microscopy slides using Fluoromount G (Thermo Fisher, cat n°00-4958-02).

### Surgeries and stereotaxic injections

Mice (postnatal day 27 - P32) were placed in a stereotaxic frame (Kopf Instrument) under isoflurane anesthesia (Baxter AG). Stereotaxic injections in the dentate gyrus (DG) region (ML 1.25 mm, AP −1.9 mm, DV −1.9 mm in C57Bl6/J background and ML 1.5 mm, AP −2.0 mm, DV −2.0 mm in BALB/C background) were performed using a Picospritzer III pressure injection system (Parker) with borosilicate glass capillaries (Hilgenberg, length 100 mm, OD 1 mm, ID 0.25 mm, wall thickness 0.375 mm). Each mouse was injected bilaterally and for each injection, 200 nL total volume was delivered through multiple spaced bursts over several minutes. After 7-9 days, mice were deeply anesthetized with ketamine/xylazine and transcardially perfused with PBS followed by PFA (4% in PBS; Electron Microscopy Sciences, cat n°15700). Extracted brains were post-fixed (PFA 4% overnight at 4°C) and stored in PBS at 4°C. Coronal brain sections were cut at 50 μm thickness with a vibratome (Leica Microsystems VT1000) and kept in PBS before staining with DAPI for 5 min and washed with PBS. Sections were subsequently mounted on glass microscopy slides with Fluoromount G (Thermo Fisher, cat n°00-4958-02).

### Immunohistochemistry

For cell-type specificity experiments, further staining was performed on the brain slices after vibratome cutting. The 50μm coronal sections were permeated in PBS 1X + Triton X 0,05% for 5 min at RT with gentle shaking and washed with PBS 1X. Blocking was performed 1h at RT in 10% Neutral Donkey Serum (Jackson ImmunoResearch, 017-000-121) and Triton-X 0,05% with gentle horizontal shaking. Primary antibody was added onto the slices and incubated overnight at 4°C with gentle shaking. The following commercial primary antibodies were used: anti-GFAP (Novus biologicals, NBP1-05198), anti-MAP2 (Synaptic systems, 188002), anti-parvalbumin (PV, Columbia University). Slices were washed with PBS 1X and incubated in secondary antibody for 1h at RT. The following commercial secondary antibodies were used: Cy5-conjugated donkey anti-chicken (Jackson ImmunoResearch, 703-175-155) and Cy5-conjugated donkey anti-mouse (Jackson ImmunoResearch, 715-175-150). Slices were again washed with PBS 1X, stained with DAPI 1/10 000 for 5 min and washed with PBS 1X. Slices were subsequently mounted on glass microscopy slides with Fluoromount G (Thermo Fisher, 00-4958-02) and imaged within one week.

### Image analysis and quantification

Confocal snapshot images and stacks were acquired from fixed HEK293T cells and brain slices with a Zeiss point scanning LSM700 confocal microscope (10X NA 0.45, 20X NA 0.80, 63X NA 1.3). For the analysis of mCherry_D4H/YDA or M/P_mCherry signal enrichment at the plasma membrane, lines (10 pixels width) were drawn across cells from single planes of 63X image stacks in Fiji ^30^. Histograms of GFP and mCherry signal intensity were derived from the lines and transferred to an Excel file. Further analysis and quantification were performed using GraphPad Prism software. To detect mCherry-positive puncta in cells, single planes of 63X image stacks were imported into Fiji, where cells were outlined manually and added to the ROI manager using the eGFP cytoplasmic filler as template. Images were then segmented to generate binary masks and particles for each cell were defined using the particle analysis function. The calculated particles were added to the ROI manager. Areas and intensities of cells and particles region were calculated for the eGFP and mCherry channels respectively and the data transferred to an Excel file. A post-hoc size cut-off of 2 pixels (0,04 μm^2^) was applied to filter particles. Further analysis and quantification were performed in GraphPad Prism software. For global qualitative assessment of in vivo injection success, images of entire coronal sections images were obtained from the Zeiss AxioScan Z1 microscope (5X NA 0.25).

### Plasma metabolites analysis

To analyse plasma metabolites, P6 or P30 *Ptchd1*^*KO*^ or wild-type male mice were deeply anesthetized using isoflurane (Baxter AG) and decapitated with scissors. Blood was collected in 300 μL Li-Heparin microvette (Sarstedt) on ice and centrifugated at 200 g for 5 min at 4°C. The supernatant (plasma) was transferred to an Eppendorf tube and stored at −20°C. Samples were diluted 1:3 in ddH_2_O on ice in a 45 μL total volume before analysis in the Cobas c111 Analyzer (Roche). The Cobas c111 Analyzer is a platform for clinical chemistry testing of human samples but it can also be used to analyze mouse samples. In this study, the following metabolites were measured: cholesterol (CHOL2), cholesterol HDL (HDLC3), cholesterol LDL (LDL-C) and triglycerides (TRIGL).

### Lipidomics

For lipid extraction, P6 or P26-28 Ptchd1 KO and wild-type male mice were deeply anesthetized using isoflurane (Baxter AG), cerebella were dissected out, flash frozen in liquid nitrogen and stored at −80°C. Cerebella were subsequently crushed into a fine powder using a custom-made metal mortar on dry ice to keep the tissue frozen. Samples were then analyzed using the following methyl-ter-buthyl ether (MTBE) modified extraction protocol. Brielfy, 5-10 mg of dry tissue were resuspended on ice in 100μL H_2_O, to which 50 μL 1.4mm Zirconium oxide glass beads (Bertin Technologies), 360 μL methanol and internal standard mix were added. The tissue was broken with 3 bursts of 45 s at 6200 rpm with 45 s interruptions in a pre-cooled Cryolysis System (Bertin Technologies) at 4°C. 1.2 mL MTBE were added before a 1h incubation in room temperature (RT) with shaking at 750 rpm. 200 μL ddH2O was added to the mixture to induce phase separation. After 10 min of incubation at RT, the samples were centrifuged at 1000 x g for 10 min. The upper organic phase was transferred to a 13 mm glass tube. The lower phase was re-extracted using 400 μL of artificial upper phase solution (MBTE/methanol/water, 10:3:1.5, v/v) and with the same incubation/centrifugation parameters. Both upper phases from extraction and re-extraction were combined and the total lipid extract was divided in 3 equal aliquots: one for phospholipid analysis (TL, total lipids one for sphingolipid analysis (SL) and one for sterol analysis (S). Aliquots were dried in a Centrivap at 50°C or under a nitrogen flow. The TL and S aliquots were ready for mass-spectrometric analysis and stored at −80°C. SL aliquot underwent further deacylation to eliminate phospholipids by methylamine treatment (Clarke method). 0.5 mL monomethylamine reagent (methanol/H_2_O/n-butanol/methylamine, 4:3:1:5, v/v) was added to the dried lipid. Samples were then sonicated and incubated for 1h at 53°C and dried as described above. The monomethylamine-treated lipids were desalted by n-butanol extraction, where 300 μL H_2_O saturated n-butanol was added to the dried lipids. The samples were vortexed, sonicated and 150 μL mass-spectrometry grade water was added. The mixture was vortexed and centrifuged at 3200 x g for 10min. The upper phase was collected and the lower phase was extracted twice more with 300 μL H_2_O saturated n-butanol. The upper phases were combined and dried as described above.

For phospholipid and sphingolipid detection, samples were pipetted into a 96-well plate with a final volume of 100 μL and LC-MS or HPLC grade solvent were used (positive mode solvent: chloroform/methanol/H_2_O 2:7:1 v/v + 5 mM ammonium acetate, negative mode solvent: chloroform/methanol 1:2 v/v + 5 mM ammonium acetate. The TL and SL aliquots were resuspended in 500 μL chloroform/methanol (1:1 v/v) solution and sonicated. The TL and SL fractions were diluted 1:10 and 1:5 respectively in a negative and positive mode solvents. For the identification and quantification of phospho- and sphingolipid molecular species, all diluted fractions were infused onto the mass spectrometer. Mass spectrometry was performed using multiple reaction monitoring (MRM) with a TSQ Vantage Triple Stage Quadrupole Mass Spectrometer (Thermo Fisher) equipped with a robotic nanoflow ion source (Nanomate HD, Advion Biosciences). Ceramide species were also quantified with a loss of water. Lipid concentrations were calculated according to the standard curves with the internal standards and normalized to the total lipid extract content. Data were also normalized between experiments. The data were not corrected for isotope distribution.

For sterol detection, the S aliquots were resuspended in 500 μL chloroform/methanol (CHROMASOLV LC-MS grade 1:1) and sonicated. 5 μL of samples were loaded onto a VARIAN CP-3800 Gas Chromatograph equipped with a Factor Four Capillary Column VF-5ms and analyzed by a VARIAN 320 mass spectrometer triple quadrupole with electron energy set to −70 eV at 250°C and the transfer line at 280°C. Temperature was held for 4 min at 45°C and ramped successively to 195°C (20°C/min), 230°C (4°C/min), 325°C (20°C/min) and 350°C (6°C/min) before cooling back to 45°C. Free sterols were eluted during the linear gradient from 195°C to 230°C. For each sterol, specific ions (m/z) were extracted: ergosterol (396.4, 363.3 and 337.3), cholesterol (386.4, 368 and 275.2) and cholesterol esters (368.4, 353.4 and 147.1). The area under the peak was integrated and the values were transferred into Excel. The sterol concentrations were determined using the standard curves of ergosterol, cholesterol and cholesterol esters.

### Statistical methods and data availability

Sample sizes conformed to 3R principles, past experience with the experiments and literature surveys. Pre-established exclusion criteria were defined to ensure success and reliability of the experiments: for stereotaxic injection, all mice with mistargeted injections were excluded from analysis (e.g. if no eGFP signal was detected in the granule cell layer of the DG). Furthermore, all mice exhibiting visible behavioral abnormalities or disease symptoms, for example in the case of the NPC1-/-mice, were excluded. Investigators performing image analysis and quantification were blinded to genotype. The applied statistical tests were chosen based on sample size, normality of data distribution and number of groups compared. Appropriate correction for variance differences was applied when necessary. Details on sample size, p-values, and specific tests are indicated in figure legends.

## Results

### Alterations in regulators of sterol metabolism are associated with neurodevelopmental conditions

Genes involved in lipid regulation are enriched amongst deleterious variants that segregate with ASD in the population ^12, 31^. To extend these observations specifically to cholesterol homeostasis, we used the GeneTrek tool ^29^ to search for neurodevelopmental disorders (NDDs) associated with mutations in genes controlling cholesterol metabolism (89 genes, KEGG pathways hsa04979 and hsa00900; Table S1). We also searched for NDD-associated genes predicted to contain sterol-sensing domains (SSD) (13 genes, Table S1) ^32, 33^, a membrane domain contained in proteins involved in cholesterol biosynthesis (e.g. HMG-CoA reductase and 7-dehydrocholesterol reductase enzymes), intracellular cholesterol transport (NPC1), and cholesterol efflux involved in cell signaling (Patched-1, PTCH1). Notably, 20 of the 89 genes involved in sterol metabolism were classified as associated with epilepsy or NDDs (**Fig. 1A**). When focusing on SSD proteins, four of the 13 genes are defined as risk genes: *NPC1, NPC1L1, PTCH1*, and *PTCHD1. NPC1, NPC1L1* and *PTCH1* encode proteins with cholesterol transporter functions. For *PTCHD1*, knock-out in mice results in behavioral and electrophysiological alterations ^27, 34, 35^. Moreover, loss of PTCHD1 impairs cholesterol-dependent μ-opioid receptor trafficking and function ^36^. However, it is unknown whether loss of *Ptchd1* affects cholesterol homeostasis *in vivo*. This close association of NDD and cholesterol metabolism genes warrants further investigation to elucidate alterations in brain cholesterol homeostasis in NDD models. Thus, we sought to develop new tools to probe cholesterol distribution in the mouse brain.

**Figure 1.**
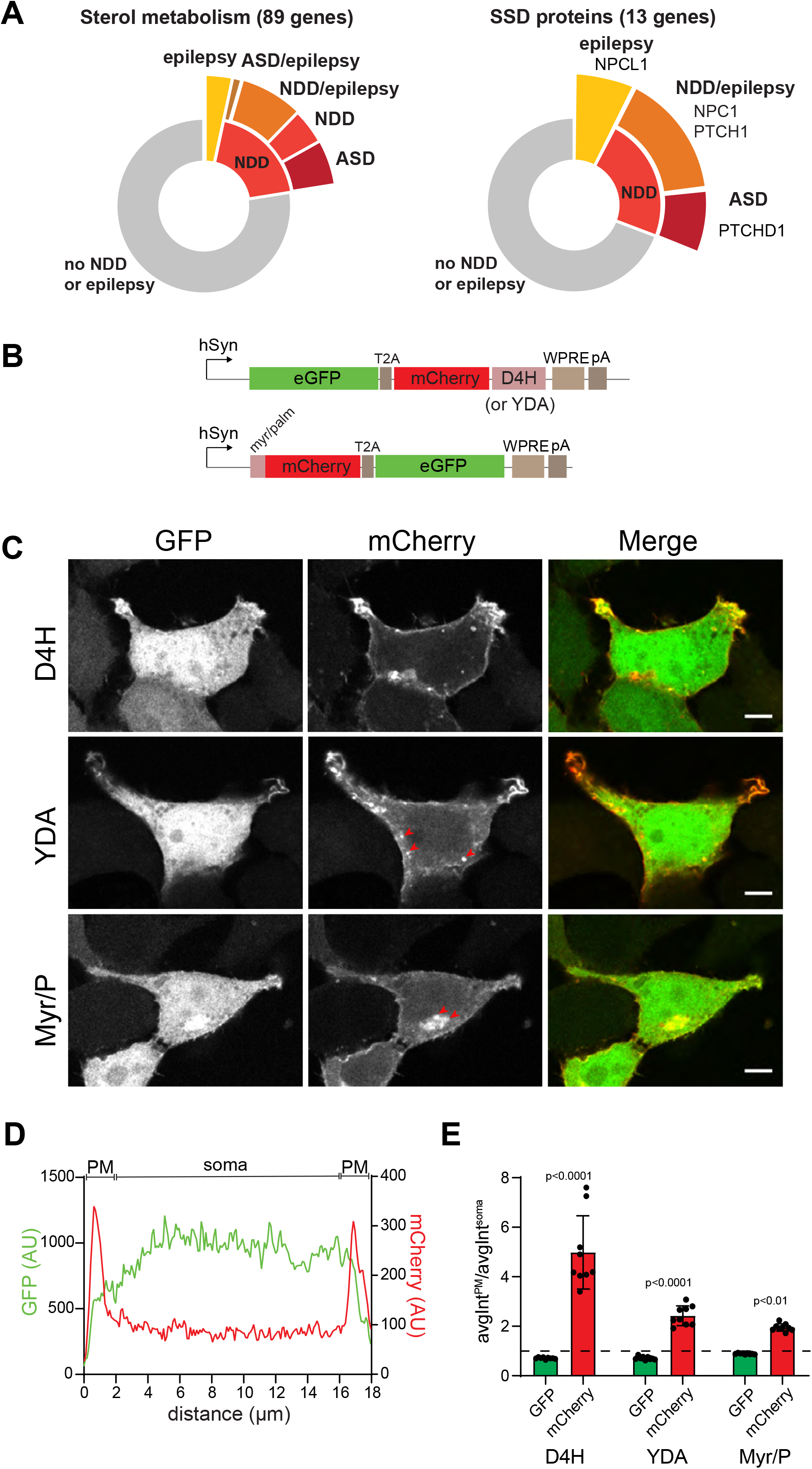
Adaptation of perfringolysin O-derived cholesterol probes for analysis of cholesterol distribution in rodent models. **A**. Pie chart results from GeneTrek analysis of curated gene lists of sterol metabolism and sterol-sensing domain (SSD) proteins (89 and 13 genes respectively). GeneTrek is used to explore association of human genes with several neurodevelopmental disorders (NDD) including autism-spectrum disorders (ASD), intellectual disability (ID) and others. Some sterol metabolism genes as well as SSD proteins genes are neurodevelopmental disorders candidate genes including *NPC1, PTCH1* and *PTCHD1*. **B**. Design of ratiometric D4H cholesterol probes under control of the human synapsin promoter. **C**. Confocal images of HEK293T cells, transfected with Syn-eGFP-2A-mCherry-D4H, Syn-eGFP-2A-mCherry-YDA or Syn-Myr/P-mCherry-2A-eGFP plasmids. Some presumptive intracellular membrane structures are marked with red arrowheads. Scale bar: 5μm. **D**. Intensity line plots of eGFP (green) and mCherry (red) signals across a single HEK293T cell. Line width used for quantification: 10 pixels (ca. 1 μm). **E**. Plasma membrane (PM) enrichment ratios of eGFP (green) and mCherry (red) for each cholesterol probe (D4H, YDA or Myr/P) plasmid in HEK293T cells. PM enrichment is calculated from average peak maximum intensity (from two consecutive pixels) at PM divided by average somatic intensity (avgInt^PM^ / avgInt^soma^). Mean ± SD, N=3 independent experiments, n=3 coverslips per condition (avg. 18 cells per coverslip). Two-way ANOVA with Tukey’s multiple comparisons tests, with individual variances computed for each comparison.

### Adaptation of cholesterol sensors for in vivo applications in mice

We aimed to measure cholesterol levels in the mouse brain in a cell type-specific manner using genetically encoded cholesterol sensors. To allow for normalization of sensor signals to expression levels, we introduced a second fluorescent protein into the construct using a T2A cleavage site for equimolar expression. Equivalent constructs were generated for the D4H and YDA sensors that are recruited to cellular membranes containing 20%mol and 2%mol cholesterol, respectively (**Fig. 1B**; D4H: eGFP-2A-mCherry-D4H and YDA: eGFP-2A-mCherry-YDA). To visualize cellular membranes independently from their cholesterol content we created a second probe targeting mCherry to membranes via a myristoylation and palmitoylation motif (Myr/Palm-mCherry-2A-eGFP, Myr/P). The probes were placed under control of the human synapsin promoter which drives neuron-specific expression in the mouse brain but also active in cultured HEK293T cells. In transiently transfected HEK293T cells, D4H signal was highly concentrated at the plasma membrane (PM) with few intracellular structures labeled (**Fig.1C**). By contrast, the YDA signal was more widely distributed consistent with labeling of intracellular membrane compartments that exhibit lower cholesterol content than the plasma membrane. As expected, the Myr/P probe labeled the PM and perinuclear membranes. Line intensity plots confirmed the intense labeling of the plasma membrane with mCherry-D4H and the presence of eGFP signal in the cytoplasm (**Fig. 1D, E**). By comparison, the YDA and Myr/P probes exhibited reduced plasma membrane enrichment due to their association with intracellular structures.

Next, we used our probes to measure cell-specific cholesterol distribution in vivo. We used stereotaxic injection of recombinant adeno-associated viruses (AAV9 capsid) to deliver the probes under control of the neuron-specific human synapsin promoter into the dentate gyrus of C57Bl6/J mice (**Fig. 2A**). Seven to nine days following injection, probes were imaged in hippocampal brain sections from perfusion fixed mice. Morphology, location of labeled cells, and co-staining with neuron and astrocyte-specific markers confirmed neuron-specificity of probe expression (**Fig. S1**). Interestingly, D4H and YDA probes strongly labeled dendrites of dentate granule cells with only low signal intensity detected in the granule cell somata (**Fig. 2B**). By contrast, cell somata located in the hilus (which mostly represent GABAergic interneurons) were strongly D4H and YDA-positive. Comparison to the eGFP normalizer as well as the Myr/P probe revealed that this was not a consequence of higher sensor expression in the hilar cells but rather a high abundance of cellular membranes within somata of these cells (**Fig. 2B**). In dentate granule cells, D4H signal was highly enriched in the plasma membrane with a small number of intracellular structures labeled. Similar to the *in vitro* conditions, YDA and Myr/P signals exhibited less plasma membrane enrichment due to the increased association with intracellular structures (**Fig. 2C**). These experiments demonstrated the suitability of D4H-derived cholesterol sensors for analysis of neuronal cholesterol distribution in the mouse hippocampus.

**Figure 2.**
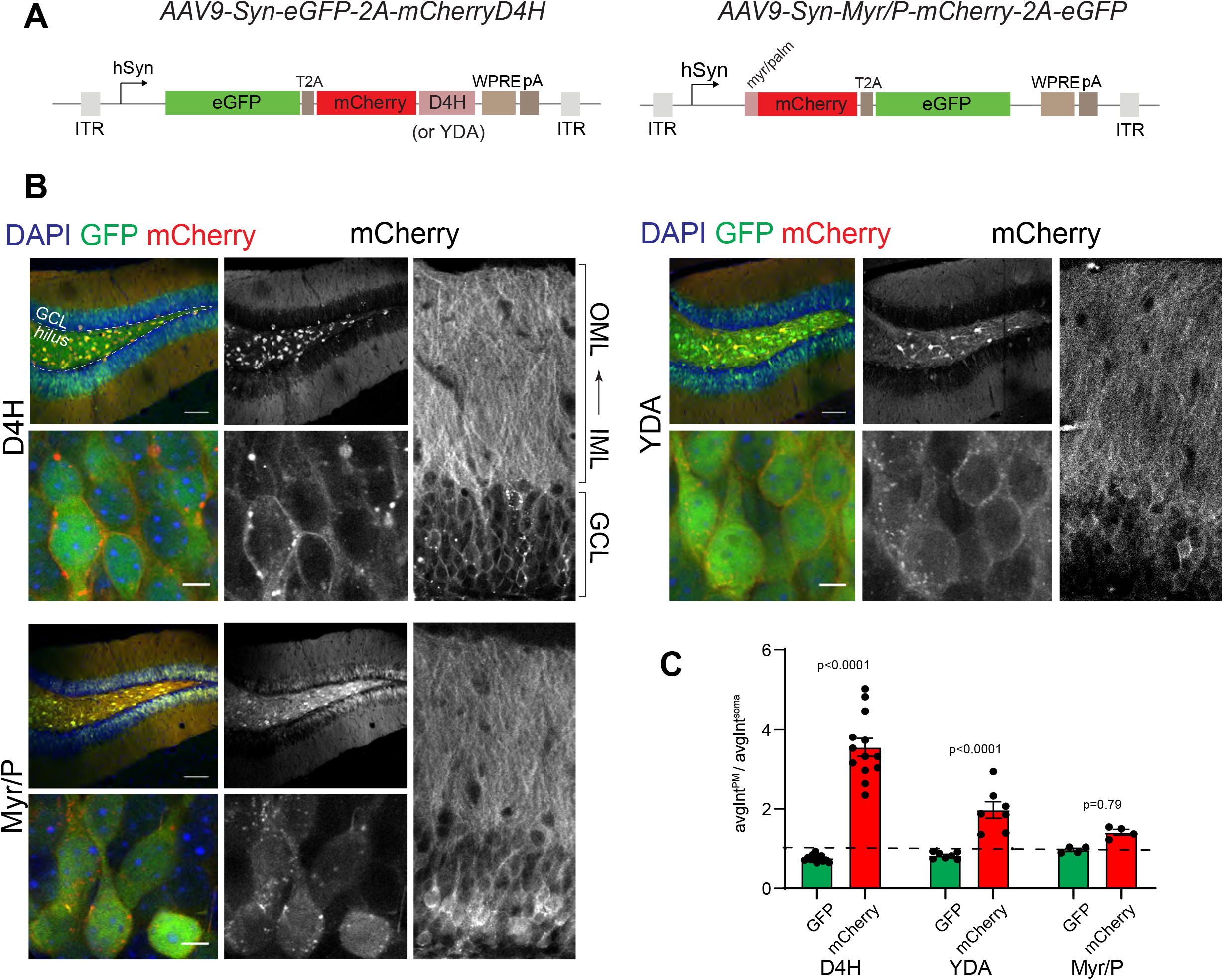
Application of ratiometric D4H cholesterol probes in mice. **A**. Design of ratiometric D4H cholesterol probes under human synapsin promoter control for neuronal expression contained in viral vectors. **B**. Confocal fluorescence micrographs of hippocampal sections from virus-injected P27-32 mice (C57B6/J background). Upper left panel: low magnification view of dentate gyrus, anatomical position of hilus and granule cell layer (GCL) is marked. Scale bar: 100μm. Lower left panel: high magnification view of granule cells, scale bar: 10 μm. Right panel: view of granule cell layer (GCL) and dendrites in inner and outer molecular layer of the dentate gyrus. **C**. Plasma membrane (PM) enrichment ratios of eGFP (green) and mCherry (red) for each cholesterol probe (D4H, YDA or Myr_P) virus in DG granule cells. PM enrichment is calculated from average peak maximum intensity (from two consecutive pixels) at PM divided by average soma intensity (avgIntPM/avgIntsoma). Mean ± SD, n=4-13 mice per condition (avg. 22 cells per mouse), Two-way ANOVA with Tukey’s multiple comparisons tests, with individual variances computed for each comparison.

### Visualization of neuron-specific cholesterol distribution in *NPC1*^*KO*^ hippocampus

Considering that mutations in the SSD protein NPC1 are associated with neurodevelopmental defects and alterations in cholesterol transport, we then used *NPC1*^*KO*^ mice to validate the suitability of the probes to quantitatively capture alterations in cholesterol distribution *in vivo*. We expressed the D4H sensor in neurons of the dentate gyrus of 4 week old wild-type and *NPC1*^*KO*^ (-/-) mice (**Fig. 3A)**. *NPC1*^*KO*^ mice displayed a strong accumulation of D4H-positive puncta in neuronal somata. These puncta resembled cholesterol accumulation in lysosomes, a cellular hallmark of Niemann-Pick Type C disease. Increased puncta density as compared to control littermates was not due to alterations in sensor expression level or cell size as confirmed by comparable eGFP signal intensity and area (**Fig. 3B**). While a small number of D4H probe-positive puncta were observed in littermate control animals, there was a substantial increase in mean puncta density and size in dentate granule cell neurons of the *NPC1*^*KO*^ mice (**Fig. 3B**). Moreover, the frequency distribution of D4H puncta density and size was strongly shifted towards higher densities and areas, with a trend towards increased probe intensity observed for these structures (**Fig. 3C**).

**Figure 3.**
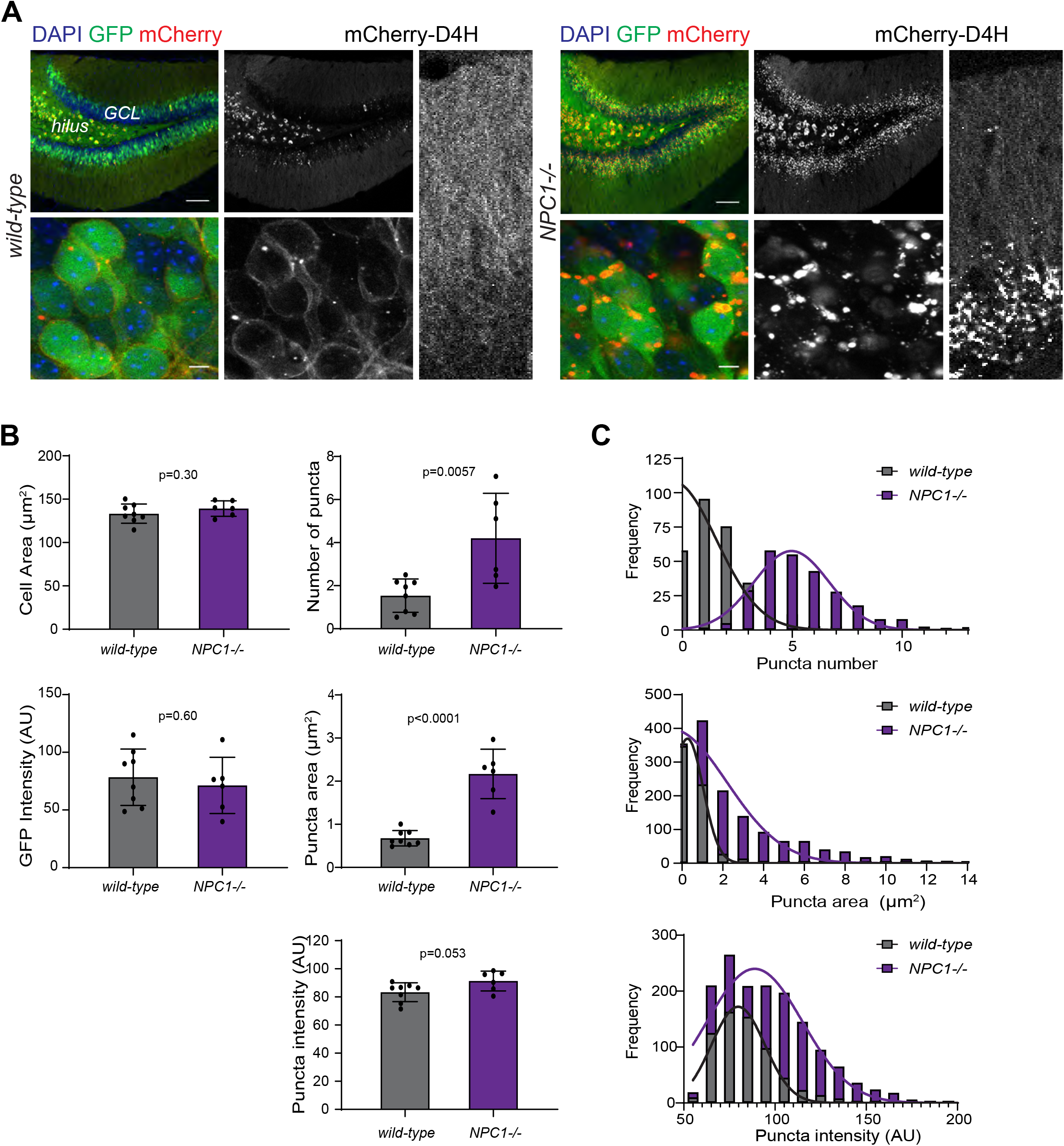
Neuronal cholesterol distribution in *Npc1*^*KO*^ mice. **A**. Confocal fluorescence micrographs of hippocampal sections from virus-injected P27-32 wild-type and knock-out (-/-) NPC1 mice. Upper left panel: Low magnification view of DG, scale bar: 100μm. Lower left panel: high magnification view of granule cells (GC), scale bar: 10μm. Right panel: low magnification view of granule cell layer (GCL) and corresponding dendrites.???again, the layers/structures are not indicated in the photos **B**. Analysis of mCherry-positive puncta in DG granule cells from NPC1 -/- mice and wildtype littermates. Left and right graphs display cell- and puncta-specific measures, respectively. Mean ± SD, n=6-8 mice per condition (avg. 162 cells per mouse), unpaired Student’s t-test. **C**. Frequency distributions of parameters (number per cell, area and intensity) with Gaussian fit.

### Normal intracellular cholesterol distribution in *Ptchd1*^*KO*^ mice

*Ptchd1* knock-out mice represent a second NDD-model carrying a mutation in a sterol-sensing domain protein. In the mouse brain, *Ptchd1* mRNA is most highly expressed in neurons of the thalamic reticular nucleus, cerebellar granule cells, and in dentate granule cells of the hippocampus ^27, 35, 37^ and genetic deletion of Ptchd1 results in defects in neuronal function in the thalamic reticular nucleus and dentate granule cells. To test whether loss of *Ptchd1* results in alterations in neuronal cholesterol distribution in granule cells, we delivered the ratiometric D4H probes into the dentate gyrus of 4-week-old male mice and compared membrane cholesterol levels between *Ptchd1* ^-/y^ mutant and littermate control animals. We detected no significant difference between genotypes in D4H probe signal in the plasma membrane or intracellular structures (**Fig. 4A-D**). This suggests that loss of *Ptchd1* from dentate granule cells is not accompanied with detectable changes in cholesterol levels or distribution in these cells.

**Figure 4.**
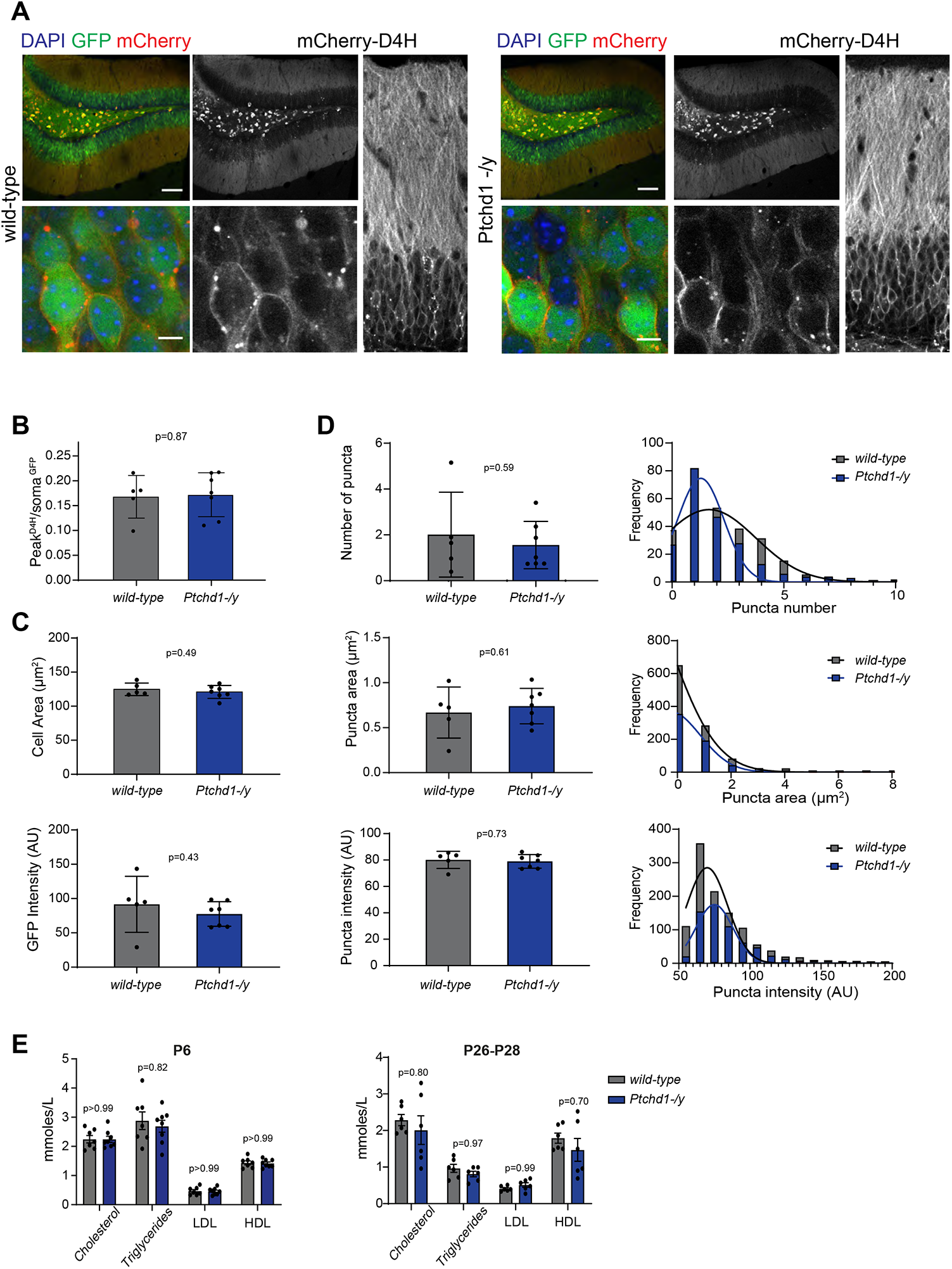
Neuronal cholesterol distribution in *Ptchd1*^*KO*^ mice. **A**. Confocal fluorescence micrographs of hippocampal sections from virus-injected P27-32 wild-type or knock-out (-/y) Ptchd1 mice (C57B6/J background). Upper left panel: Low magnification view of DG, scale bar: 100 μm. Lower left panel: high magnification view of granule cells (GC), scale bar: 10 μm. Right panel: view of granule cell layer (GCL) and corresponding dendrites. **B**. Plasma membrane (PM) enrichment ratios of eGFP (green) and mCherry (red) in DG granule cells of injected wild-type and *Ptchd1*^*KO*^ mice. PM enrichment is calculated from average peak maximum intensity (from two consecutive pixels) at PM divided by average soma intensity (avgInt^PM^/avgInt^soma^). Mean ± SD, n=10-13 mice per condition (avg. 45 cells per mouse), multiple unpaired Student’s t-test with Holm-Sidak method for statistical significance determination. **C**. Analysis of mCherry puncta in DG granule cells from *Ptchd1* wild-type and *Ptchd1*^*KO*^ mice. Each cell was manually traced and mCherry puncta were identified by intensity thresholding. Left and right graphs display cell- and puncta-related parameters, respectively. Mean ± SD, n=5-7 mice per condition (avg. 166 cells per mouse), unpaired Student’s t-test. **D**. Frequency distributions of puncta-related parameters with Gaussian fit. **E**. Plasma concentration of cholesterol-related metabolites from P6 or P30 wild-type and *Ptchd1*^*KO*^ mice. LDL: low density lipoprotein, HDL: high density lipoprotein. Mean ± SD, n=7-8 mice, two-way ANOVA with Tukey’s multiple comparisons tests, with individual variances computed for each comparison.

Considering that *Ptchd1* is also expressed in non-neuronal tissues ^37^ and that alterations in blood cholesterol levels were reported in ASD patients ^12, 13^, we performed blood sampling from 6 day and 30 day old *Ptchd1*^*-/y*^ and littermate control animals and quantified cholesterol, LDL, HDL, and triglycerides levels. Notably, we did not observe significant genotype differences in any of these parameters at both ages (**Fig. 4E**). Finally, no significant changes in overall lipid composition, including fatty acid chain length or saturation were observed using mass-spectrometry-based lipidomic analysis of *Ptchd* ^-/y^ mutant brain samples (Fig. S2 and Supplementary Table 2). Taken together, these results suggest that deletion of the sterol-sensing domain protein PTCHD1 does not significantly alter cholesterol level or other lipid levels in the mouse brain.

### Selective targeting of sensors reveals cell type-specific differences in cholesterol distribution

A major advantage of molecularly-encoded sensors is the ability to target them to genetically-defined cell populations. Considering the segregation of brain cholesterol synthesis and cholesterol turn-over between astrocytes and neuronal cells, respectively, we modified the AAV vectors to target the probes to other cell types. Thus, we expressed the ratiometric D4H probe selectively in astrocytes of the dentate gyrus using a minimal GFAP promoter (*GfaABC1D)* (**Fig.5A**) ^28^. Cholesterol distribution in astrocytes *in vivo* differed significantly from neuronal cells, with an accumulation of probe-positive structures in the astrocytic cytoplasm in wild-type mice (**Fig. 5B**). Interestingly, these astrocytic cholesterol-containing structures were reduced in number and only slightly enlarged in *Npc1*^*KO*^ mice (**Fig.5C**). This highlights a highly cell type-specific phenotype of NPC1 loss-of-function in astrocytes where the main cholesterol depots arise from new synthesis as opposed to neurons. To further expand the range of cell populations that can be targeted with ratiometric D4H probes in mice, we generated viral vectors for Cre recombinase-dependent expression. We validated those vectors by selective expression in parvalbumin-positive interneurons in the mouse dentate gyrus (**Fig. S3**). In combination, these versatile tools should prove highly valuable to complement lipidomic and molecular studies providing cell type and sub-cellular resolution for assessment of cholesterol distribution *in vivo*.

**Figure 5.**
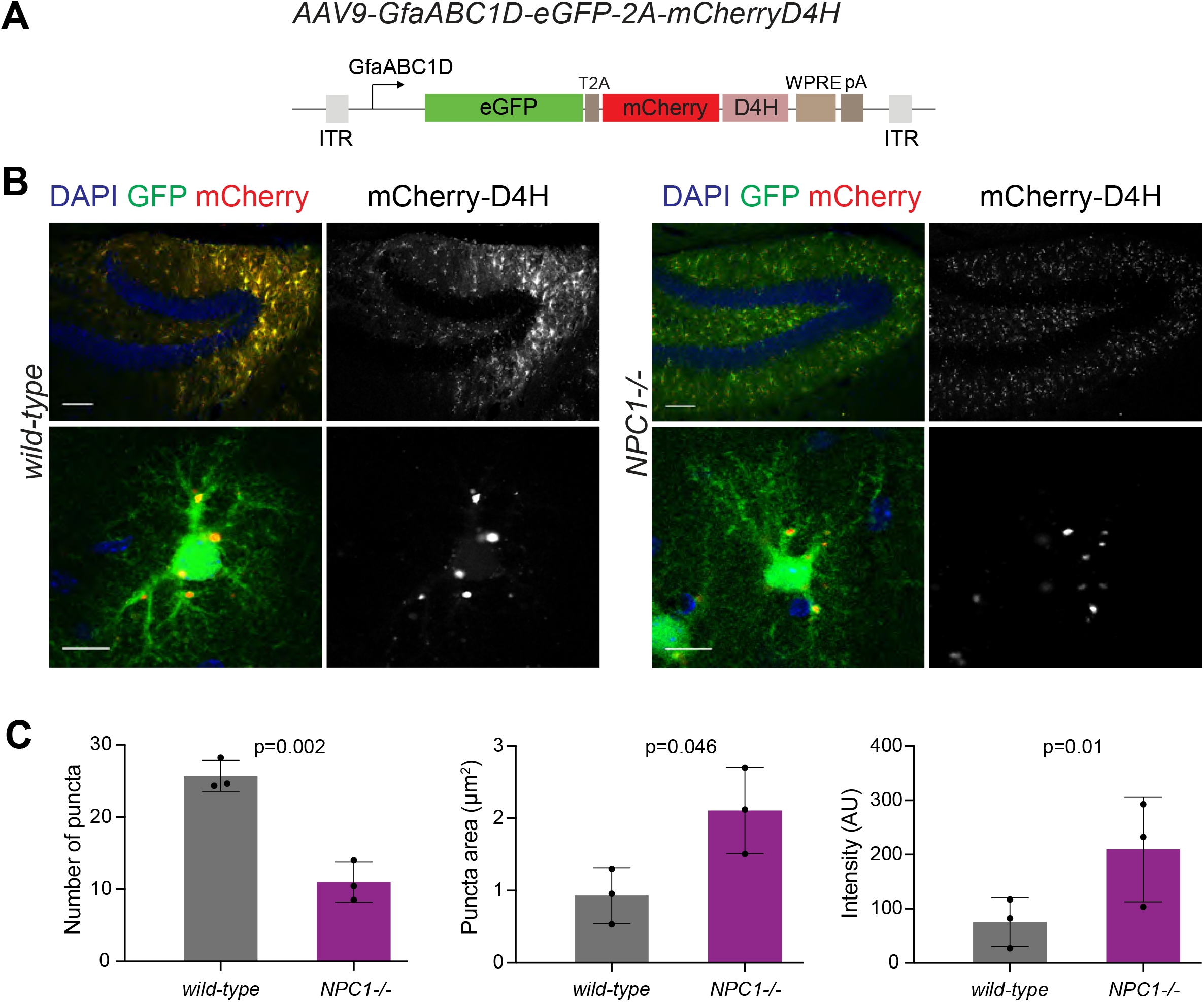
Adaptation of D4H probes for probing astrocyte-specific cholesterol distribution. **A**. Expression cassette for viral expression of eGFP-T2A-mCherryD4H cholesterol probe under control of the astrocyte-specific GfaABC1D promoter. **B**. Confocal flluorescence micrographs of hippocampal sections from P27-32 wild-type or *Npc1*^*KO*^ mice after stereotaxic injection of AAV9 GfaABC1D vectors driving probe expression. Top panels: Low magnification view of dentate gyrus, scale bar: 100 μm. Lower panels: High magnification view of probe-expressing of astrocyte, scale bar: 10 μm. **C**. Analysis of mCherry-positive puncta in DG astrocytes from wild-type and *NPC1*^*KO*^ mice. Mean ± SD, n=3 mice per condition (total of 42 cells from wild-type animals and 35 cells from *NPC1*^*KO*^ animals), unpaired Student’s t-test.

## Discussion

Mounting evidence has linked lipid – and more particularly cholesterol – metabolism alterations to neurodevelopmental disorders. Neuronal cholesterol must be tightly controlled as either increased or decreased levels are associated with disease states. As a key component of cellular membrane and membrane microdomains, cholesterol modulates ion channel function, signaling receptors, and synaptic vesicle release ^38-40^. For example, cholesterol is required for function of multiple GPCRs ^41, 42^, including the high-affinity state of the oxytocin receptor, a receptor centrally involved in the regulation of social behaviors ^43^. Consequently, disrupted cholesterol homeostasis has severe impact on brain function ^44, 45^. A deeper understanding of cell type-specific alterations in cholesterol homeostasis associated with disease states might open up new therapeutic avenues, particularly, as cholesterol-modulating drugs are available.

D4-derived cholesterol sensors have been employed previously to map distribution of pools of cholesterol in cultured cells ^24, 26, 46, 47^ and in *C*.*elegans* ^48^. Here, we adapted these probes for cell type-specific measurements in the rodent brain. A key modification was the introduction of a second fluorophore that enables ratiometric measurements and introduction of a cholesterol-independent membrane probe. This combination of probes allows for normalization to probe expression and differences in intracellular membrane content across cell types. We demonstrate the utility of these probes to capture cholesterol distribution across brain cell types and to visualize cell type-specific cholesterol trafficking defects in the *NPC1*^*KO*^ model. Interestingly, we observed highly divergent cholesterol distribution phenotypes in *NPC1*^*KO*^ astrocytes and granule cells in the mouse dentate gyrus. While loss of NPC1 resulted in accumulation of cholesterol in granule neurons, the probe-accessible pools in hippocampal astrocytes were only modestly affected. This observation is consistent with the proposal that anatomically-defined phenotypes can be rescued by neuron-specific re-expression of NPC1 protein ^49^. The cre recombinase-dependent sensors developed here should facilitate the dissection of cholesterol trafficking defects in additional cell types in the *NPC1*^*KO*^ model, like microglia, where impaired lysosomal lipid trafficking, enhanced phagocytic uptake, and impaired myelin turnover have been reported ^50^.

In contrast to *NPC1*^*KO*^ mice, we did not detect significant alterations in the D4H-accessible cholesterol pool in *Ptchd1*^*KO*^ animals. Moreover, overall sterol levels, lipid composition or acyl chain saturation in *Ptchd1* mutants were unchanged. This suggests that PTCHD1 protein is unlikely to have a major cholesterol transport function in the mouse brain. While NPC1 and PTCHD1 share sterol-sensing domains, they differ in key amino acid residues linked to transport function in the RND permease protein family. NPC1 and the sonic hedgehog receptor PTCH1, two SSD proteins with cholesterol transport function contain GxxxDD and GxxxE/D motifs in transmembrane domains 4 and 10, respectively ^51, 52^. Notably, in PTCHD1, these motifs are not conserved, similar to SCAP another SSD protein (**Fig. S4**). SCAP is involved in cholesterol biosynthesis regulation where it acts as cholesterol sensor ^53^. Notably, a recent study linked PTCHD1 and plasma membrane cholesterol to μ-opioid receptor trafficking ^36^. Thus, it might be of interest to further investigate sterol-sensing and cholesterol-dependent trafficking functions of PTCHD1 in the future.

A subset of individuals with autism have characteristic dyslipidemia including cholesterol alterations, which can be identified by analysis of blood samples ^12, 13^. Initial trials in individuals with Fragile X have not supported therapeutic benefits from treatment with cholesterol lowering drugs. The approach developed in our work should enable probing brain cholesterol distribution on a cellular level in models of brain disorders or in patient-derived cell preparations. Such an analysis should provide valuable information on forms of ASD, where a modification of cholesterol levels may hold promise for therapeutic intervention. Finally, alterations in cholesterol homeostasis are more widely implicated in brain pathology, including Alzheimer’s Disease ^54^. Thus, the virally-delivered probes developed in our work should accelerate capturing cell type-specific cholesterol distribution in the mammalian brain and enable tracking alterations in cholesterol flow across cell types in a wide-array of brain disorders.

## Acknowledgements

We are grateful to Drs. Ozgur Genc and Anne Spang for insightful comments on the manuscript, Dr. Sophie Martin for sharing cDNAs of the cholesterol probes, and colleagues in the Scheiffele lab for support and advice. This work received funding from AIMS-2-TRIALS which are supported by the Innovative Medicines Initiatives from the European Commission. The results leading to this publication has received funding from the Innovative Medicines Initiative 2 Joint Undertaking under grant agreement No 777394. This Joint Undertaking receives support from the European Union‘s Horizon 2020 research and innovation programme and EFPIA and AUTISM SPEAKS, Autistica, SFARI. Moreover, the work has received funding from the European Union‘s Horizon 2020 research and innovation programme under grant agreement No 847818 (CANDY) and NCCR SYNAPSY. The funders had no role in the design of the study; in the collection, analyses, or interpretation of data; in the writing of the manuscript, or in the decision to publish the results.

