## Supplementary Figures for "Cell type-specific assessment of cholesterol distribution in models of neurodevelopmental disorders"

**Extended Data**

**Inventory of Supplementary Information**

**Table S1.** List of genes encoding proteins involved in cholesterol metabolism and proteins containing sterol-sensing domains.

**Table S2.** Detailed results of lipidomic analysis.

**Supplementary Figure 1.** Data confirming neuron-specific expression of synapsin promoter-driven cholesterol probes

**Supplementary Figure 2.** Additional data on lipidomic comparison of wild-type and *Ptchd1*$^{KO}$ tissue.

**Supplementary Figure 3.** Additional data on cell type-specificity of cre-dependent cholesterol probe expression.

**Supplementary Figure 4.** Amino acid sequence motifs related to a potential transporter function of sterol-sensing domain proteins.

**Legends related to Tables.**

**Table S1.** List of genes encoding proteins involved in cholesterol metabolism and proteins containing sterol-sensing domains.

**Table S2.** Detailed results of lipidomic analysis. Results from 5 male *Ptchd1*$^{KO}$ and male littermate wild-type mice. Abbreviations are: Cer: ceramide; GlcCer: glucosylceramide; SM: sphingomyelin; PC: phosphatidylcholine; PE: phosphatidylethanolamine; PI: phosphoinositol; PS: phosphatidylserine; CL:cardiolipin. Lengths of acyl chains are indicated in numbers (_10; _12 etc).

**Supplementary Figures**

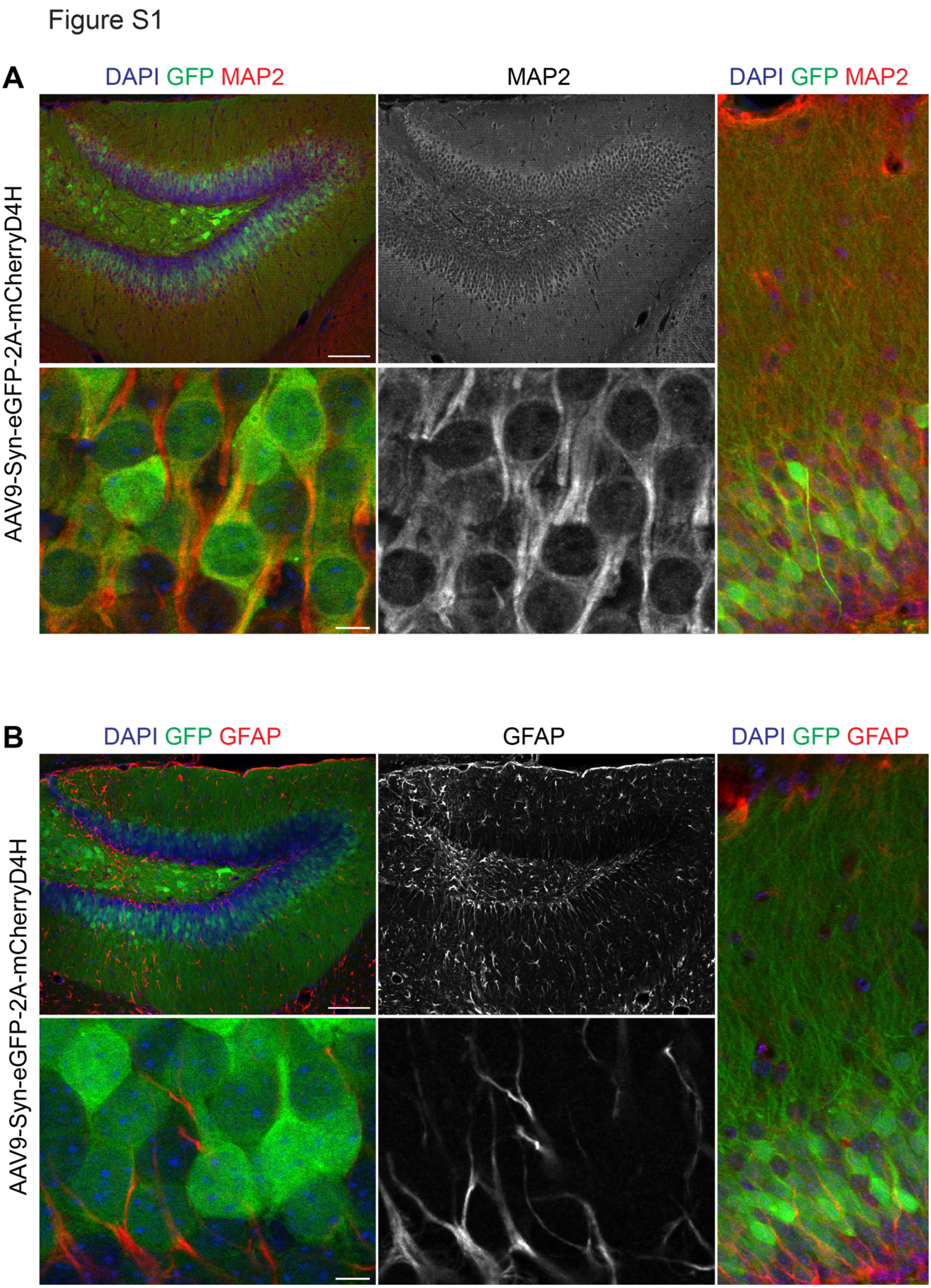

**Figure S1. Neuron-specific expression of ratiometric cholesterol probes in mice driven by the human synapsin promoter**

**A.** Confocal fluorescence micrographs showing D4H cholesterol probes expressed from AAV vectors delivered by stereotaxic injection into the adult mouse dentate gyrus. Probe-expressing cells are marked by eGFP expression (displayed in green). Neuronal cells are identified by immunostaining for the neuronal microtubule-associated protein MAP2 (displayed in red). **B.** D4H probe expression as in A with co-labelling for the astrocyte-specific glial acidic fibrillary protein (GFAP displayed in red). Upper left panels: Overview of DG, scale bar: 100 µm. Lower left panels: Higher magnification view of granule cells (GC), scale bar: 10µm. Right panel: 20X magnification view of granule cell layer (GCL) and corresponding dendrites.

### Figure S2

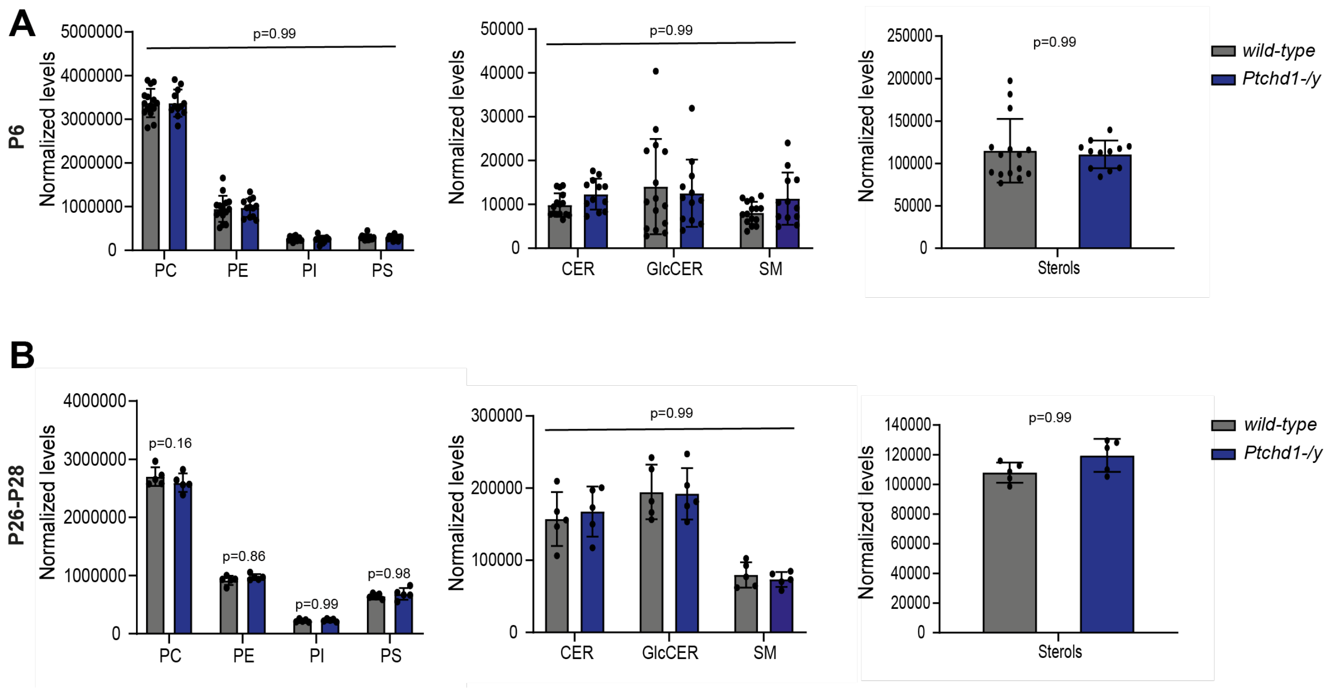


**Figure S2. Summary of lipidomic comparison of wild-type and *Ptchd1*$^{KO}$ tissue**

**A.** Lipids levels (total picomoles, normalized) in cerebellar tissue of P6 *Ptchd1* wild-type or -/y mice (n=15-12) by lipidomics analysis. **B.** Lipids levels (picomoles, normalized) in cerebellar tissue of P26-28 *Ptchd1* wild-type or -/y mice (n=5) by lipidomics analysis. Note that cerebellar tissue was used in these experiments as it is a site of high PTCHD1 mRNA expression [27]. PC: phosphatidylcholine, PE: phosphatidylethanolamine, PI: phosphatidylinositol, PS: phosphatidylserine, Cer: ceramide, GlcCer: glucosylceramide, SM: sphingomyelin. Mean ± SD, Ordinary two-way ANOVA with Tukey's multiple comparisons tests, with individual variances computed for each comparison.

Figure S3

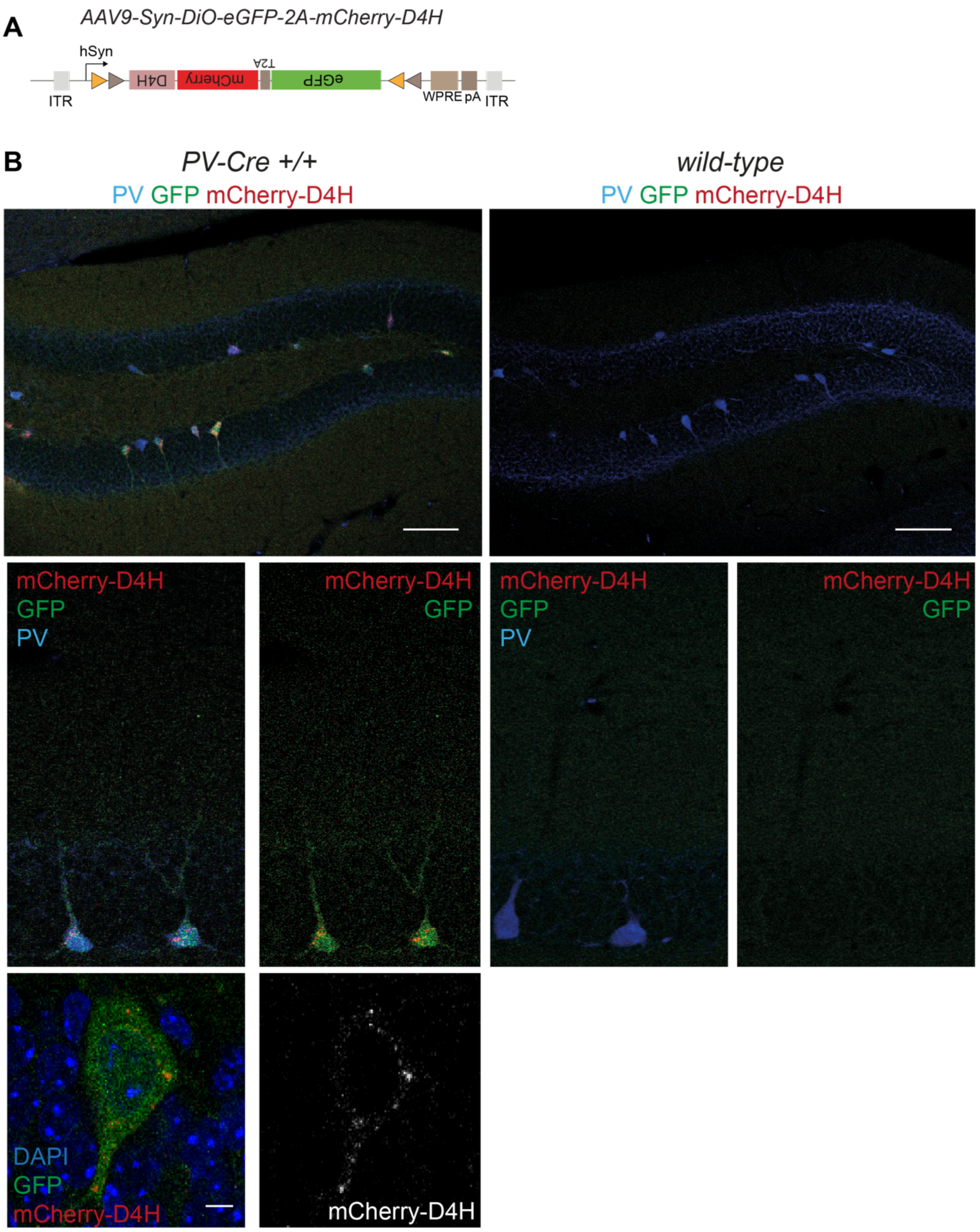


**Figure S3. AAV vectors for cell type-specific visualization of cholesterol distribution**

**A.** Design of ratiometric D4H cholesterol probe with DiO configuration for cre-dependent expression. **B.** Confocal fluorescence micrographs of hippocampal coronal sections of P27-32 wild-type or PV-Cre mice. Parvalbumin cells are identified by immunostaining of the PV protein (displayed in blue). Upper left panel: Overview of DG, scale bar: 100 µm. Lower left panel: Higher magnification view of granule cells (GC), scale bar: 10µm. Right panel: 20X magnification view of granule cell layer (GCL) and corresponding dendrites.

#### Figure S4

**A**

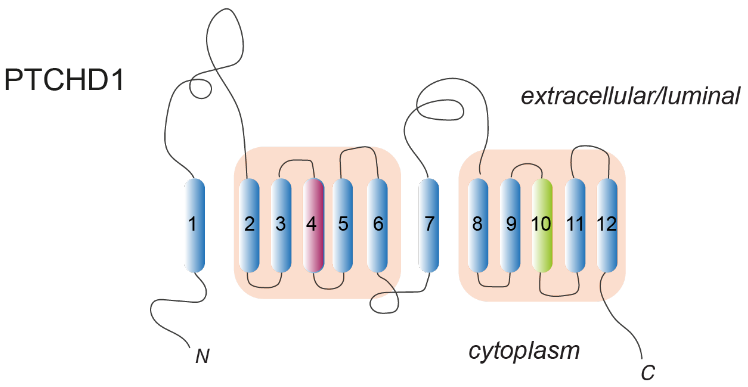


**B**

```
                 TM4
PTCHD1  TFLGVPFVMLGHGLYGTFEMLSSW
PTCH1   TTQVLPFLALGVGVDDVFLLAHAF
NPC1    VIEVIPFLVLAVGVDNIFILVQTY
DISP1   MNLTALIILVGIGADDAFVLCDVW
SCAP    GGEIFPYLVVVIGLENVLVLTKSV
                  GXXXDD
```

RND transporter
key residues

```
                 TM10
PTCHD1  KNALEVHGVAILQSYLCYIVGLFP
PTCH1   PVVILIASVGIGVEFTVHVALAFL
NPC1    SLVNLVMSCGISVEFCSHITRAFT
DISP1   ESVTISVAVGLSVDFAVHYGVAYR
SCAP    ------------------------
                 GXXXE/D
```

RND transporter
key residues

**Figure S4. Key amino acid residues of RND transporter family are not conserved in PTCHD1.**

**A.** Hypothetical topology of PTCHD1. The two predicted sterol-sensing domains are highlighted in orange, transmembrane domains in blue (numbered from 1-12). **B.** Alignment of amino acid sequences (mouse proteins) in transmembrane domains 4 and 10 which play key roles in cholesterol transport by PTCH1. Critical amino acid residues contributing to transport function in bacterial RND transporters as well as PTCH1 are highlighted (GxxxDD and GxxxD/E motif) [51,52]. NPC1 and DISP1 which also exhibit cholesterol transport activity share the conserved motifs. By contrast, SCAP – which lacks cholesterol transporter activity lacks these motifs. Note that SCAP contains only one SSD and, thus, only 8 transmembrane domains.
